# Multiomics data collection, visualization, and utilization for guiding metabolic engineering

**DOI:** 10.1101/2020.10.15.341909

**Authors:** Somtirtha Roy, Tijana Radivojevic, Mark Forrer, Jose Manuel Marti, Vamshi Jonnalagadda, Tyler Backman, William Morrell, Hector Plahar, Joonhoon Kim, Nathan Hillson, Hector Garcia Martin

## Abstract

Biology has changed radically in the past two decades, growing from a purely descriptive science into also a design science. The availability of tools that enable the precise modification of cells, as well as the ability to collect large amounts of multimodal data, open the possibility of sophisticated bioengineering to produce fuels, specialty and commodity chemicals, materials, and other renewable bioproducts. However, despite new tools and exponentially increasing data volumes, synthetic biology cannot yet fulfill its true potential due to our inability to predict the behavior of biological systems. Here, we present a set of tools that, combined, provide the ability to store, visualize and leverage these data to predict the outcome of bioengineering efforts.

## Introduction

Synthetic biology represents another step in the development of biology as an engineering discipline. The application of engineering principles such as standardized genetic parts (Canton et al., 2008; Müller and Arndt, 2012) or the application of Design-Build-Test-Learn (DBTL) cycles (Petzold et al., 2015; Nielsen and Keasling, 2016) has transformed genetic and metabolic engineering in significant ways. Armed with this new engineering framework, synthetic biology is creating products to tackle societal problems in ways that only biology can enable. Synthetic biology, for example, is being leveraged to produce renewable biofuels to combat climate change (Beller et al., 2015; Chubukov et al., 2016; Peralta-Yahya et al., 2012), improve crop yields (Roell and Zurbriggen, 2020), combat the spread of diseases (Kyrou et al., 2018), synthesize medical drugs (Ajikumar et al., 2010; Paddon and Keasling, 2014), biomaterials (Bryksin et al., 2014), and plant-based foods (Meat-free outsells beef, 2019).

However, the development of synthetic biology is hindered by our inability to predict the results of engineering outcomes. DNA synthesis and CRISPR-based genetic editing (Doudna and Charpentier, 2014; Ma et al., 2012) allow us to produce and change DNA (the working code of the cell) with unparalleled ease, but we can rarely predict how that modified DNA will impact cell behavior (Gardner, 2013). As a consequence, it is not possible to design a cell to fit a desired specification: e.g., have the cell produce *X* grams of a specified biofuel or an anticancer agent. Hence, metabolic engineering is often mired in trial-and-error approaches that result in very long development times (Hodgman and Jewett, 2012). In this context, machine learning has recently appeared as a powerful tool that can provide the predictive power that bioengineering needs to be effective and impactful (Carbonell et al., 2019; Radivojević et al., 2020; Zhang et al., 2020).

Furthermore, although there is a growing abundance of phenotyping data, the tools to systematically leverage these data to improve predictive power are lacking. For example, transcriptomics data has a doubling time of seven months (Stephens et al., 2015), and high-throughput techniques for proteomics (Chen et al., 2019) and metabolomics (Fuhrer and Zamboni, 2015) are becoming increasingly available. Often, metabolic engineers struggle to synthesize this data deluge into precise actionable items (e.g. down regulate this gene and knock out this transcription factor) to obtain their desired goal (e.g. increase productivity to commercially viable levels).

Here, we showcase how to combine the following existing tools to leverage omics data and suggest next steps (Fig. 1): the Inventory of Composable Elements (ICE), the Experiment Data Depot (EDD), and the Automated Recommendation Tool (ART). ICE (Ham et al., 2012) is an open source repository platform for managing information about DNA parts and plasmids, proteins, microbial host strains, and plant seeds. EDD (Morrell et al., 2017) is an open source online repository of experimental data and metadata. ART (Radivojević et al., 2020; Zhang et al., 2020) is a library that leverages machine learning for synthetic biology purposes, providing predictive models and recommendations for the next set of experiments. When combined, this set of tools can effectively store, visualize, and leverage synthetic biology data to enable predictive bioengineering and effective actionable items for the next DBTL cycle. We will demonstrate this with an example in which we leverage multiomics data to improve the production of isoprenol, a potential biofuel (Kang et al., 2019).

**Figure 1:**
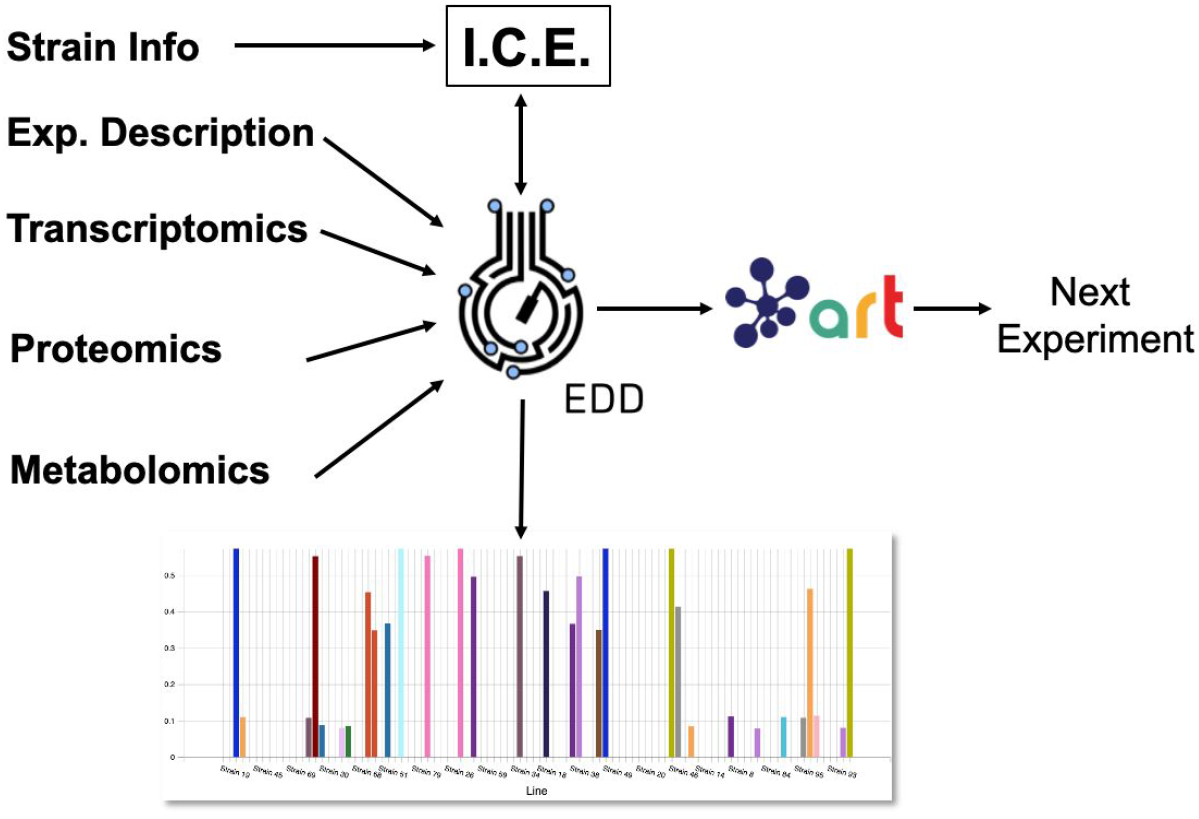
Combining several tools to guide metabolic engineering. The combination of ICE, EDD, and ART provides the ability to store, visualize and leverage multiomics data to guide bioengineering. Here, we showcase how to use this collection of tools to improve the production of isoprenol in *E. coli* for a simulated data set.

## Results and discussion

In the following sections, we will provide a step-by-step example of how to use this suite of tools (ICE, EDD, and ART) to guide metabolic engineering efforts and increase isoprenol production in *E. coli* for a synthetic data set (Fig. 2, and “Methods” section). Using synthetic (simulated) data allows for a more effective demonstration, since there is no obstacle to creating time series multiomics data involving transcriptomics, proteomics, metabolomics, and fluxomics. For the purposes of this demonstration, creating such a data set using real experiments would be prohibitively expensive and limiting.

**Figure 2:**
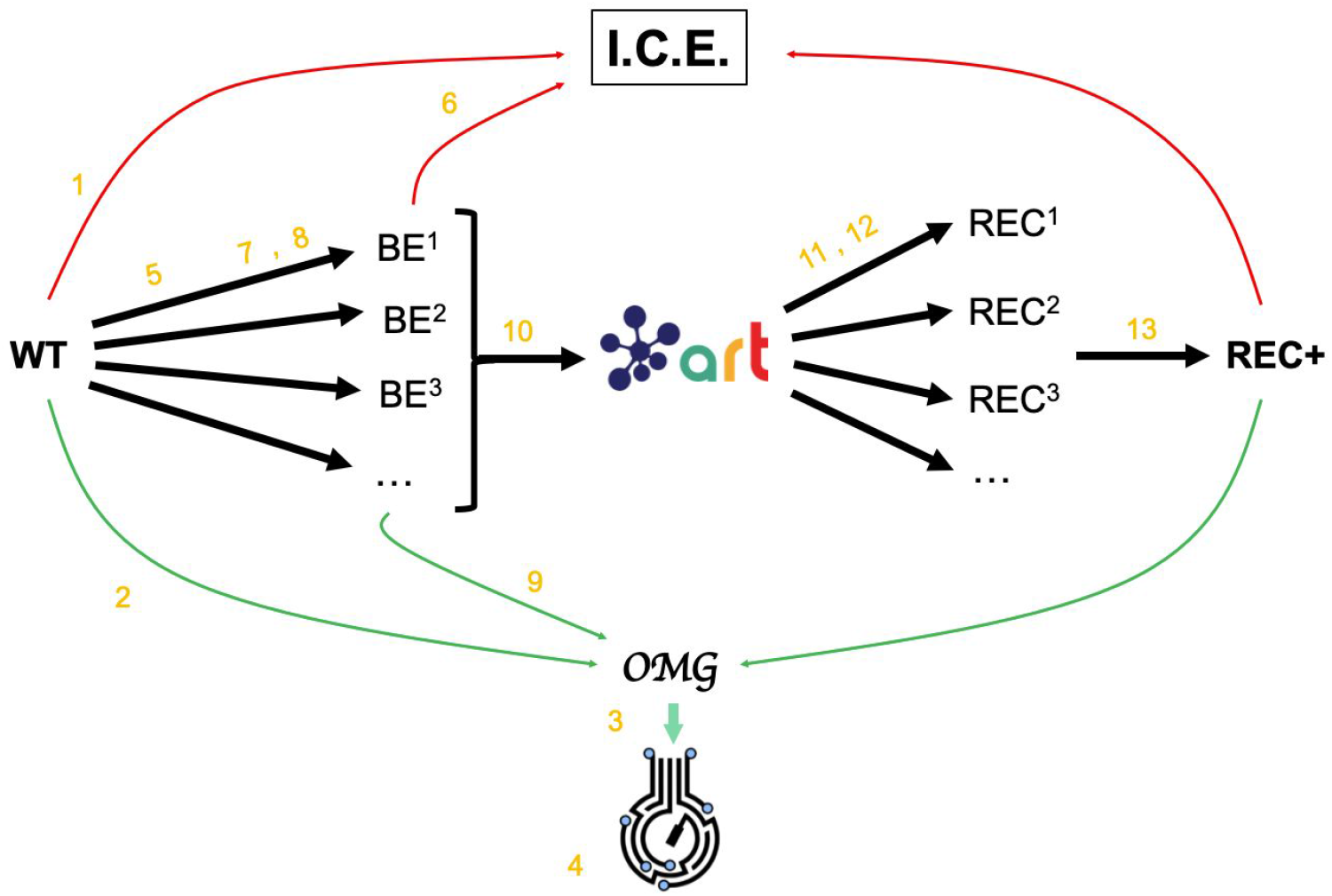
Demonstrating ICE, EDD, and ART using synthetic data. For the purposes of the demonstration of how ICE, EDD, and ART work together, we use a synthetic data set of multiomics data (transcriptomics, proteomics, metabolomics, fluxomics) for several time points created by the Omics Mock Generator (OMG) library (see Methods section). We start with a base strain (wild type, or WT) that is bioengineered according to several designs (i.e.: knockout malate dehydrogenase, overexpress citrate synthase) suggested by ART. The results are 95 bioengineered strains (BE*,BE^2^… etc) for which experimental data (isoprenol production levels) are simulated through OMG and stored in EDD and ICE. These data are then leveraged by ART to recommend, using machine learning, new designs that are expected to improve isoprenol production (REC^1^, REC^2^,…). These recommendations and production predictions are compared with the ground truth provided by OMG. Each of these steps (in orange) is demonstrated through screencasts and Jupyter notebooks (Table 1).

### Storing strain information in ICE

Our first step involves storing the initial strain information in the Agile BioFoundry (ABF) instance of ICE: https://public-registry.agilebiofoundry.org/ (see Screencast 1 in Tables 1 and 2). Storing strain information in ICE provides a standard way to document the design phase and make this information available for later use. ICE provides access controls so that the strains can be created as the experiment progresses, and then made public later (e.g. at publication time). We will initially store the information for the base strain (or wild type, WT). After creating an account and logging in, we click on “Create Entry” and choose “Strain”. We then fill the relevant strain information (e.g. “Name”, “Biosafety Level”, “Description”, “Sequence”, etc). Finally, we will click on “Submit” to create the strain entry. The strain is now available on the ICE instance and is assigned a part number that will be used to enter experimental data into EDD as the next step. Strain information can also be easily downloaded from ICE through the GUI (Screencast 5).

**Table 1:** Workflow for the paper and demonstrative jupyter notebooks and screencasts. All screencasts and notebooks are enumerated in Tables 2 and 3.

| Step | Description | Demonstration |
| --- | --- | --- |
| 1 | WT strain import into ICE | Screencast 1 |
| 2 | WT multiomics data generation | Notebook A |
| 3 | EDD import of WT multiomics data | Screencast 2 |
| 4 | EDD visualization of WT multiomics data | Screencast 3 |
| 5 | Initial designs generation by ART | Notebook B |
| 6 | Bulk ICE import of bioengineered (BE) strains | Screencast 4 |
| 7 | ICE export of BE strains | Screencast 5 |
| 8 | Isoprenol production data generation for BE strains | Notebook C |
| 9 | Bulk EDD import and visualization of BE strains production data | Screencast 6 |
| 10 | EDD export of BE production data | Screencast 7 + Notebook D |
| 11 | ART predictions and recommendations | Screencast 8 + Notebook D |
| 12 | Using the ART frontend | Screencast 9 |
| 13 | Comparing ART predictions with ground truth | Notebook E |

**Table 2:** Screencasts.

| Screencast | Description |
| --- | --- |
| 1 | WT Strain import into ICE |
| 2 | EDD import of WT multiomics data |
| 3 | EDD visualization of WT multiomics data |
| 4 | Bulk ICE import of bioengineered (BE) strains |
| 5 | ICE export of BE strains |
| 6 | Bulk EDD import and visualization of BE strains production data |
| 7 | EDD export of BE production data |
| 8 | ART predictions and recommendations |
| 9 | Using the ART frontend |

### Importing Data into EDD

Importing the multiomics data set into EDD allows for standardized data storage and retrieval. The use of standardized schemas is fundamental for downstream analysis such as machine learning or mechanistic modeling. Indeed, it is estimated that 50-80% of a data scientist’s time is spent with the type of data wrangling that EDD avoids by providing an ontology (For Big-Data Scientists, ‘Janitor Work’ Is Key Hurdle to Insights - The New York Times). This ontology reflects the objects and processes most often encountered in metabolic engineering experiments (see Fig. 2 in (Morrell et al., 2017)). For example, a typical metabolic engineering project starts by obtaining several *strains* from a strain repository (e.g. DMZ, ATCC, ICE). Those strains are cultured in different flasks under different conditions (different media, induction levels etc.), which we call *lines* because they are a combination of strain and culture conditions, and represent a different line of enquiry or question being asked (e.g. does this strain under this condition improve production?). The final steps usually involve making measurements relevant to the experiment’s ultimate goal. These measurements could be the concentration of the metabolite that the strains are engineered to produce (e.g. isoprenol in our example), or a transcriptomics or proteomics analysis that describes the amount of gene transcription or protein expression. Each of these measurements is obtained by applying a *protocol* (e.g. proteomics) to a given line, resulting in an *assay*.

Performing an assay results in a set of *measurement data*: e.g., the number of grams of acetate per liter in the media, or the number of proteins per cell. In this ontology, assays can include one or more time points. As a general rule, destructive assays (e.g. proteomics through LC-MS) include one time point per assay, and non-destructive assays (e.g. continuous measurement of cell optical density through an optode in a fermentation platform such as a biolector) include several time points in the same assay. All this information is collected in a *study*, which is used to describe a single continuous experiment (e.g. using measuring isoprenol production for all bioengineered strains).

The first step in the data import involves creating a study and uploading an “Experiment description” file, which collects all the experimental design and metadata (see Screencast 2). The “Experiment description” file describes the strains being used through a “Part ID” number tied to a strain repository, such as ICE (Fig. 3). This file also contains metadata relevant to the experiment (e.g. temperature, culture shaking speed, culture volume etc.). The “Experiment description” file should not include any result data. Often, the distinction between data and metadata is crystal clear, but it can be blurred in the case of concatenated experiments: e.g., the hydrolysate sugar concentration for a plant deconstructed with ionic liquid can be data for a deconstruction experiment, but metadata for an experiment focused on the fermentation of that hydrolysate through a bioengineered strain. As a rule of thumb, metadata involves the information that is known before the experiment, and data is the information that is only obtained by performing the experiment. Once the “Experiment description” file is added, you should be able to see all the experimental design data under the “Experiment description” tab in EDD.

**Figure 3:**
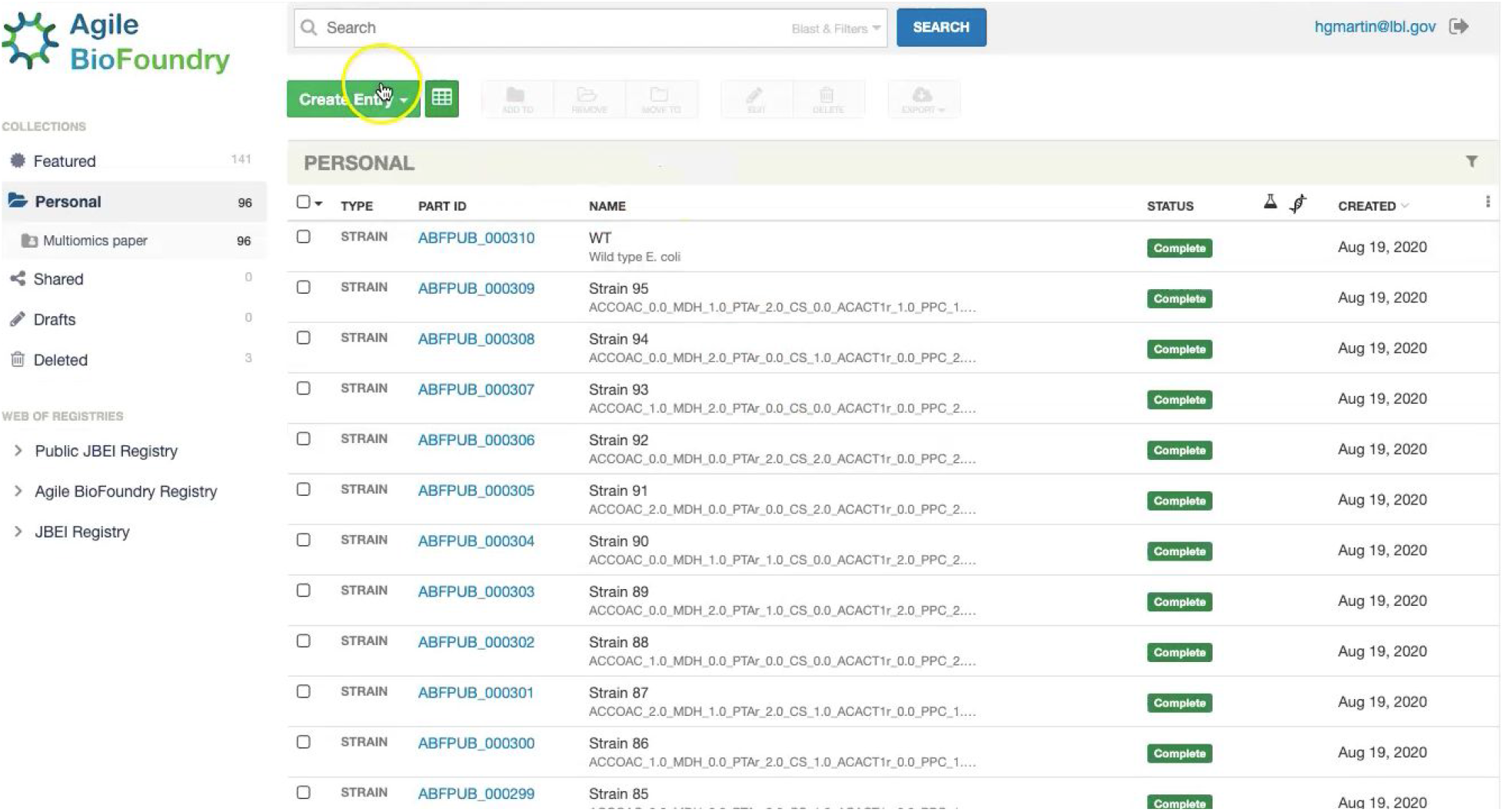
Storing strain information in ICE. ICE provides a standardized repository to store information for DNA parts and plasmids, proteins, microbial host strains, and plant seeds. These data will be linked to the experimental data contained in EDD through the part ID, to be present in the experiment description file.

The second step in data input involves uploading the data files for each assay. In order to do so, click on “Import data”, and you will access EDD’s new streamlined import. This new import emphasizes clarity and usability, and starts by prompting for the “Data category” to be uploaded (Fig. 4). Data categories involve broad umbrellas of data types such as: transcripts, proteins, metabolites, or other data. The next choice involves the actual specific protocol used for acquiring the data. There are many types of, e.g., proteomics protocols which differ in extraction protocols, as well as the type of mass spectrometer used and its setting (chromatography column, gradient time, etc). We encourage the use of formal protocol repositories which provide DOIs (digital object identifiers), such as protocols.io (Teytelman et al., 2016), to encourage reproducibility. Protocols can be added by the system administrator in charge of the EDD instance used. The next choice involves the type of file used to input the data. All protocols include a *generic* file type which is the simplest possibility: a data identifier and a single number (see “Data formats for generic input files” section below). More complex file types can be easily added through scripts that map into this generic file type. The file upload completes the data import, and these data can be now found under the “Data” tab in the study. Similar to entering strains in ICE, data and metadata import into EDD creates incremental documentation of the experiment in a form that can easily be published later by modifying the study’s access controls.

**Figure 4:**
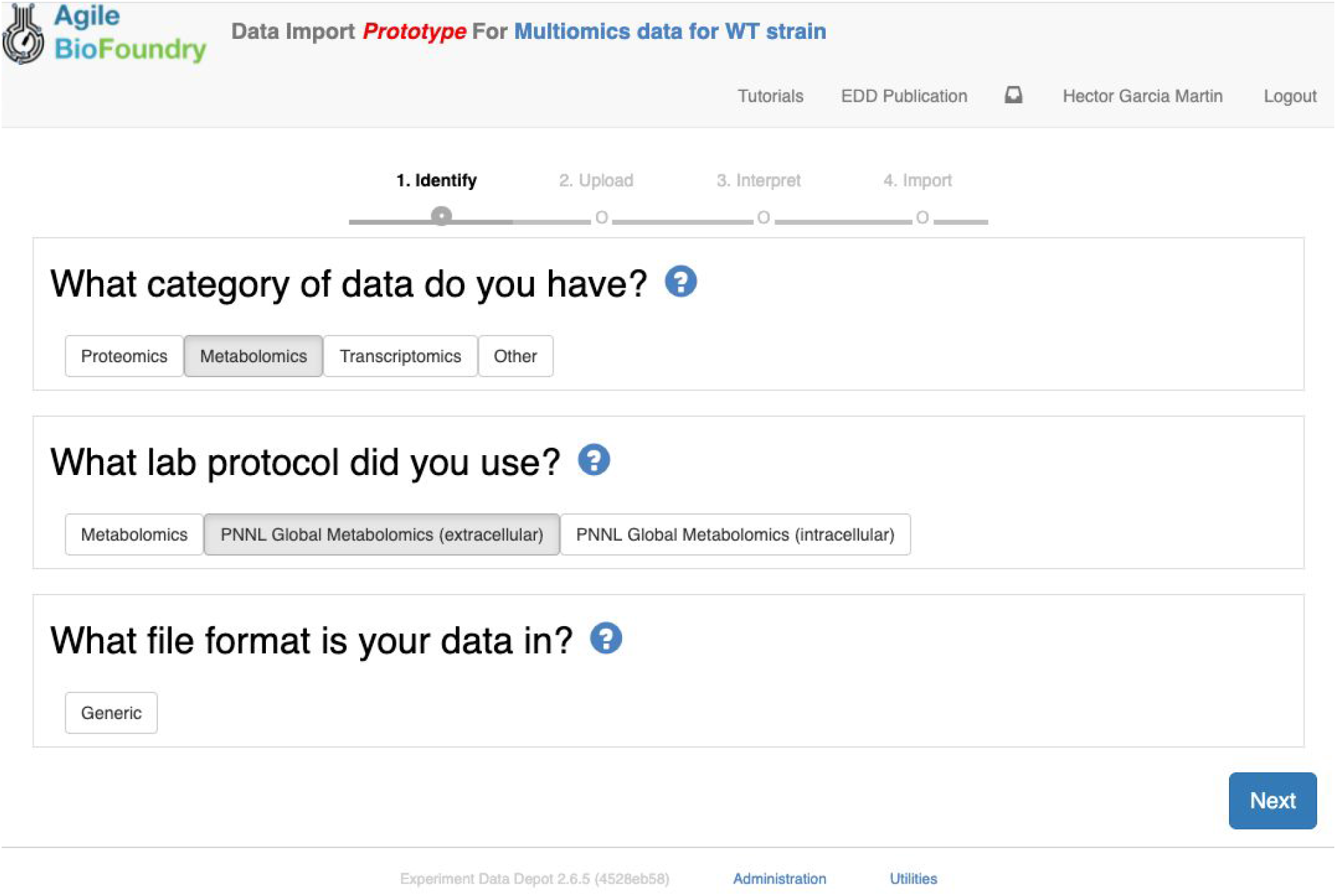
Importing data into EDD. The new data import into EDD is divided into three parts: an initial choice of the data category, the protocol used to gather the data, and the file format used for the data. Once these are chosen, the data is uploaded for future visualization and use with, e.g., machine learning algorithms or mechanistic models.

We import data into EDD twice in our metabolic engineering workflow (Fig. 2): steps 3 and 9 in Table 1. In the first case (Screencast 2), we upload data created through the OMG library for the wild type (WT, Notebook A). Later (step 8) we use OMG to simulate isoprenol production data for the 95 bioengineered strains proposed by ART (Notebook C), and upload that data into EDD (step 9, Screencast 6).

### Visualizing data in EDD

EDD provides data visualization for the comparison of multiomics data sets though line and bar graphs. For example, you can easily compare the synthetic data sets created for this manuscript, which include the cell density, extracellular metabolites, transcriptomics, proteomics and metabolomics data for the base strain (Fig. 2). Once in EDD, the data can be viewed by clicking on the “Data” tab in the corresponding EDD study (see Screencast 3, Fig. 5). The default view is the “Line Graph” view, which displays the data as times series, with the time dimension on the *x*-axis and the measurements on the *y*-axis. Each different measurement unit is given an axis (e.g., mg/L for metabolites, proteins/cell for proteomics, FPKM for transcriptomics). The filters at the bottom of the screen allow the users to choose the data they want to concentrate on: e.g., only transcriptomics, only proteomics, only metabolites, only metabolite “octanoate”, or only proteins “Galactokinase” and “Maltoporin”, or only gene “b0344” and protein “Enolase”. It is also possible to view the data in the form of bar graphs clustered by measurement, line, or time. The “Table” tab shows a quick summary of all data available in the study.

**Figure 5:**
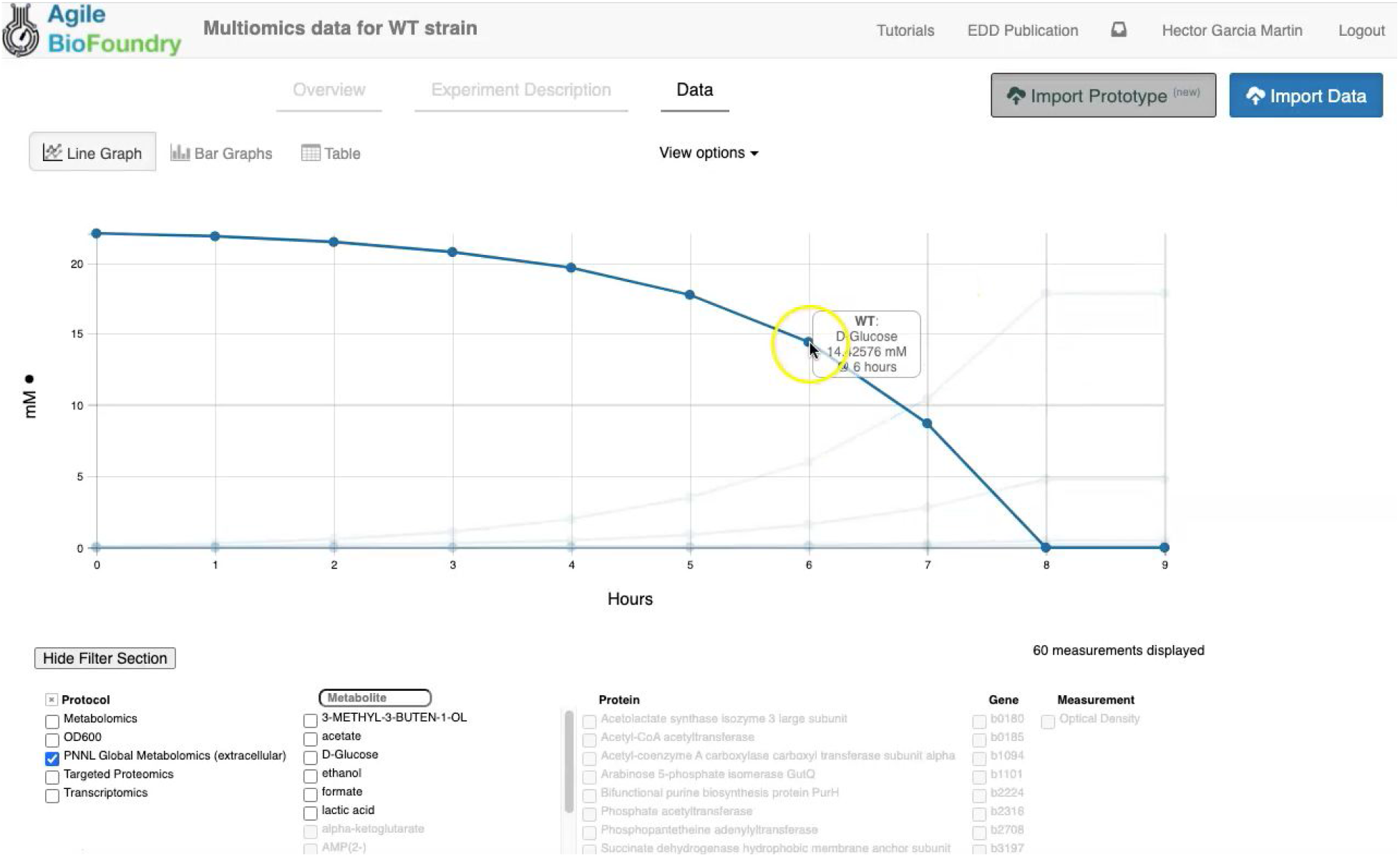
Visualizing data in EDD. EDD provides data visualization in the form of bar and line charts. The lower menu provides filtering options to facilitate comparison of lines. More sophisticated visualization can be achieved by pulling the data from EDD through the REST API.

EDD provides basic visualization capabilities to check data quality and compare different lines and data types. Users who want a customized visualization or figure can download the data into a pandas data frame (pandas - Python Data Analysis Library) through the REST API (see next section), and use Python to leverage any of the available visualization libraries: e.g., Matplotlib (Yim et al., 2018) or seaborn (seaborn: statistical data visualization — seaborn 0.10.1 documentation).

EDD visualization is demonstrated for two very different cases in two steps of our metabolic engineering workflow (Fig. 2): steps 4 (Screencast 3) and 9 (Screencast 6). The first case involves visualizing multiomics data for a single strain (WT, Fig. 5), whereas the second case involves the visualization of shallow data (final isoprenol production at a final point) for 96 different strains.

### Exporting data from EDD

Data can be exported from EDD in two ways: a manual CSV file download, or a REpresentational State Transfer (REST, (Masse, 2011)) Application Programming Interface (API). The REST API is the preferred method since it is easy to use, convenient, and flexible.

CSV export works through the Graphical User Interface (GUI) found in the “Table” tab. By selecting the desired measurements and clicking on “Export Data”, the user can access a menu that provides options for layout and metadata to be included, as well as a visual example of the export. A CSV file is then generated by clicking on “Download”.

The REST API provides a way to download the data in a form that can be easily integrated into a Jupyter notebook (see Fig. 6, Screencast 7, and Table 3). A Jupyter notebook is a document that contains live code, equations, visualizations and explanatory text (Project Jupyter | Home; IOS Press Ebooks - Jupyter Notebooks - a publishing format for reproducible computational workflows). The edd-utils package uses EDD’s REST API to provide a DataFrame inside of your Jupyter notebook to visualize and manipulate as desired. A DataFrame is a structure from the popular library Pandas (Python ANd Data AnalysiS, (pandas - Python Data Analysis Library)), that focuses on providing tools for data analysis in python. The combination of Pandas with Jupyter notebooks provides reproducible workflows and the capability to do automated data analyses (see Screencast 8). The users can run the `export_study ()` function from the edd_utils package (see code availability) to download a study from a particular EDD instance. The study is identified by its slug: the last part of the internet address corresponding to the EDD study (Fig. 6). The EDD instance is identified by its internet address (e.g., http://public-edd.agilebiofoundry.org/ or https://public-edd.jbei.org/). Anyone with an approved account can use the EDD instances hosted at those addresses, or anyone can create their own EDD instance by downloading and installing the open source EDD software (see software availability). EDD includes access controls, e.g. for preventing dissemination of experimental data prior to publication.

**Figure 6:**
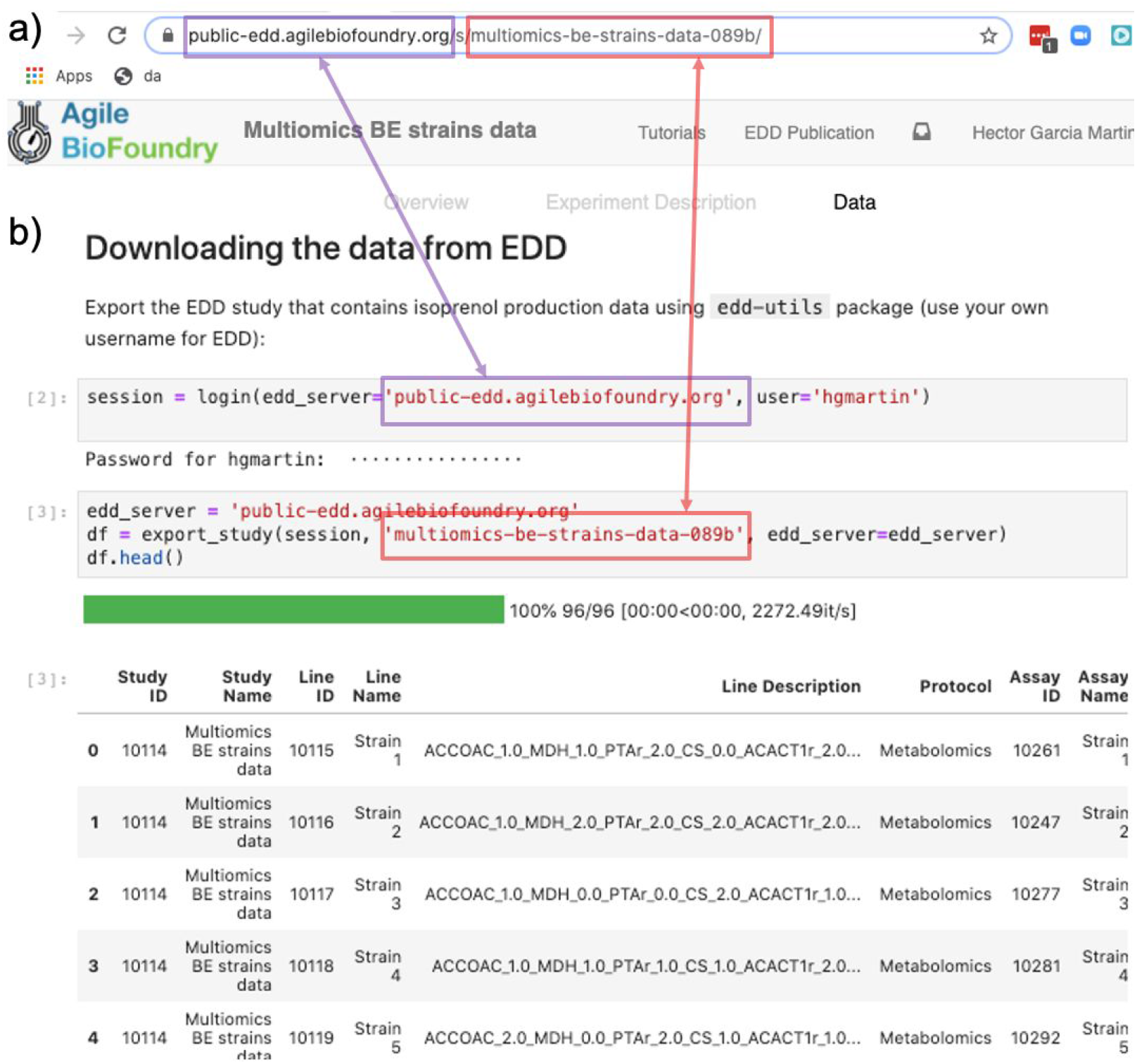
Exporting data from EDD into an executable Jupyter notebook for downstream processing. The EDD study web address (**a**) provides the server (magenta) and the slug (red) to export the study data in the form of a pandas data frame into a Jupyter notebook (**b**). Once in a data frame format in a Jupyter notebook, a plethora of Python libraries are available for visualization, mechanistic modeling or machine learning.

**Table 3:**
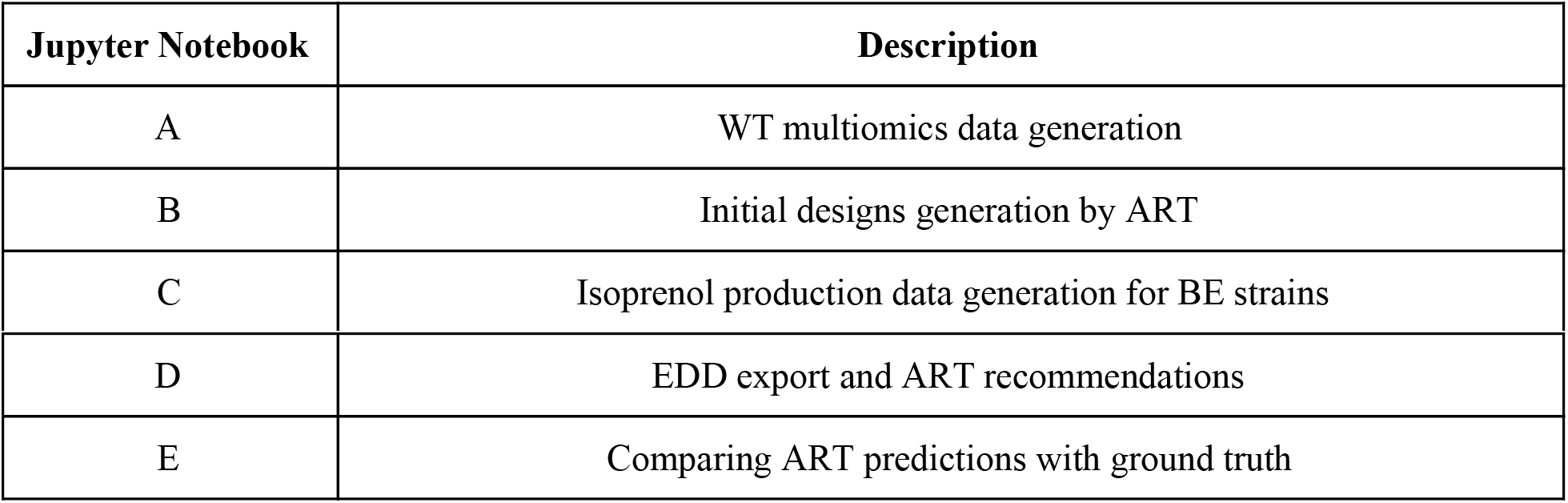
Jupyter notebooks.

EDD export is demonstrated in a single step in our workflow (Fig. 2): the export of production data for the bioengineered strains in step 10 (Screencast 7). These data will be used to train the machine learning algorithms in the Automated Recommendation Tools, and recommend new designs.

### Recommending new designs through ART

The data stored in EDD and ICE can be used to train machine learning methods and recommend new experiments. We will now show, for example, how to use the data we uploaded into EDD to suggest how to improve the final production of isoprenol, by using the Automated Recommendation Tool (ART, (Radivojević et al., 2020)). ART is a tool that combines machine learning and Bayesian inference to provide a probabilistic predictive model of production, as well as recommendations for next steps (Fig. 7). In this case, we will use as input the genetic modifications on the strain (e.g., knockout ACCOAC, overexpress MDH, maintain CS, etc.) and we will try to predict final isoprenol production (see “Generating training data for machine learning and testing predictions” section).

**Figure 7:**
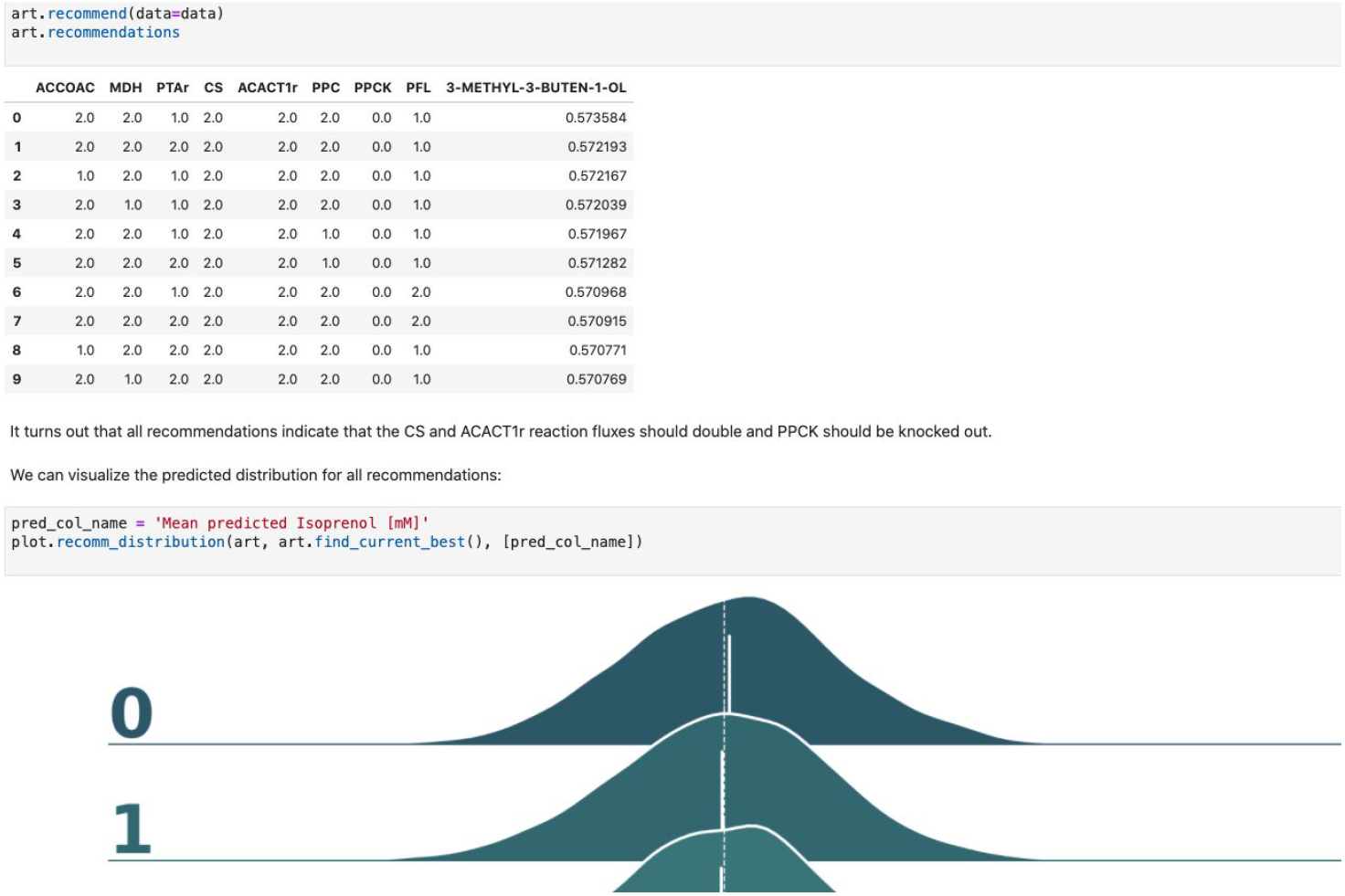
Using machine learning to predict production and recommend new designs. The ART library takes a DataFrame containing input designs (i.e. which fluxes to overexpress, 2, keep the same, 1, or knock out, 0) and isoprenol production (response). The trained model recommends new designs that have the highest production. The recommendations come with predictions of production in a probabilistic fashion: i.e. the probability of production 10, 15, 25, 40 mMol, etc.

Firstly, we will adapt the DataFrame obtained previously (step 10 in Table 1) to provide training data for ART. All we need to provide ART is the input (genetic modifications) and response (final isoprenol production) for each of the 96 instances. We do this by expanding the line description to include the design details into several new columns detailing the specified genetic modification for each reaction (Screencast 8, Notebook D). Next, we specify ART’s parameters: input variables, response variable, number of recommendations, etc. ART uses these data and parameters to train a predictive model, which is able to recapitulate quite effectively the observed isoprenol production for the training data set (Fig. 8). Recommendations for designs that are predicted to increase production are also provided by ART (Fig. 7).

**Figure 8:**
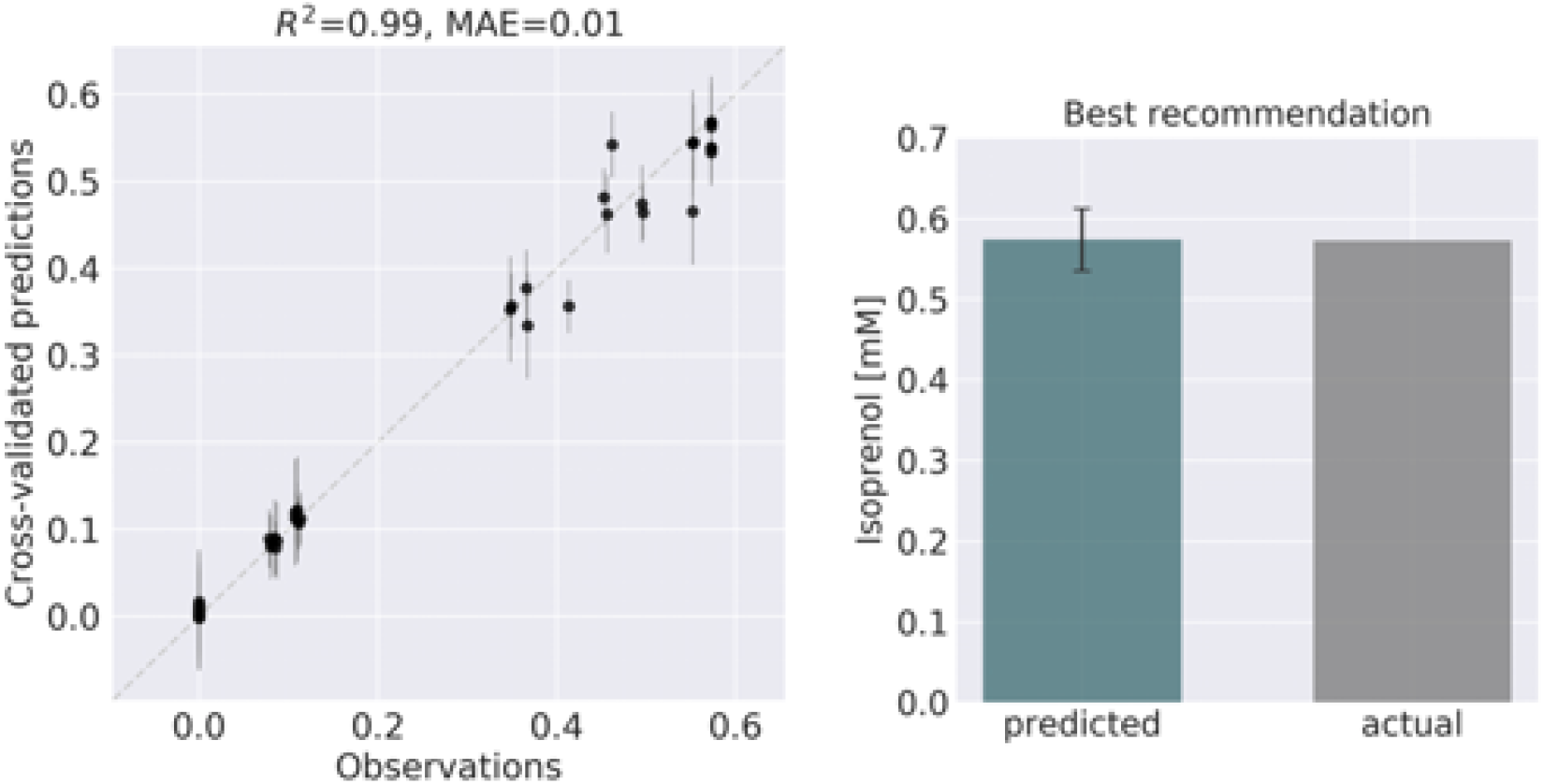
ART recommendations display production levels production very similar to predictions. Left panel compares cross-validated predictions for isoprenol production from ART versus the values obtained through the OMG library for the training data set. Cross validation keeps a part of the data set hidden from the training to compare against predictions, providing a good idea of the quality of predictions for new data sets. The right panel compares the predicted production for the recommended strain (#97) vs the actual production as generated through the OMG library. The comparison indicates a very good agreement between the prediction and observation.

We can also use ART’s web-based graphical frontend to produce recommendations, if we prefer not to use code. ART’s frontend can be found at https://art.agilebiofoundry.org/, and requires creating an account (Screencast 9). Once the account is created and approved, we can provide the same DataFrame we created above in CSV or Excel format (Notebook D). We can then select the input variables and response variables, select the objective (maximization, minimization or reach a target value), a threshold for declaring success, and then press “Start”. The job is sent to the server and an email is sent to the user once it is finished.

ART’s recommendations suggest that knocking out the PPCK flux and either maintaining or overexpressing all other seven fluxes should increase production of isoprenol from 0.46 mMol to 0.57 mMol (23 % increase, Fig. 7). The machine learning model suggests that these are the best combinations based on the predictive probability. We can check this result through the OMG library, which we consider our ground truth (Fig. 2). Indeed, isoprenol production for this design is 0.57 mMol vs the predicted 0.57 ±0.02 (Fig. 8). Hence, ART has been able to predict which combination of designs would produce a production increase. This is a non-trivial endeavour, since only 11% of designs actually improve production, according to the synthetic data provided by OMG.

**Figure 9:**
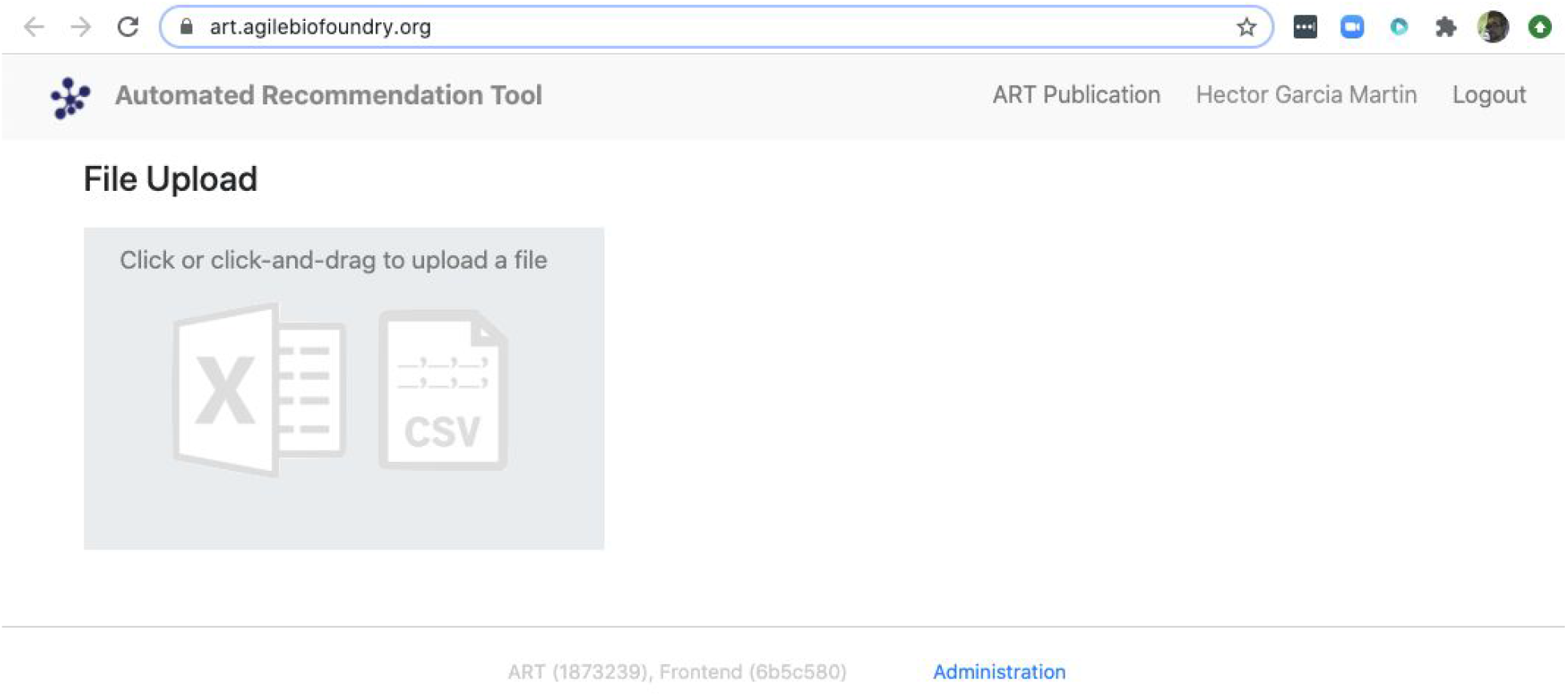
ART also provides a frontend that does not require coding. The frontend can be found at https://art.agilebiofoundry.org/ and provides the main functionality of the ART library (Fig. 7) in an intuitive interface. The frontend also provides a REST API that users with coding experience can leverage to use Berkeley Lab’s compute resources for running ART, or to trigger ART runs automatically from other code.

## Conclusion

In conclusion, we have shown that the combination of tools presented here (ICE + EDD + ART + Jupyter notebooks) provide a standardized manner to store data so it can be leveraged to produce actionable recommendations. We have shown how to use ICE to store strain information, EDD to store experiment data and metadata, and ART to leverage these data to suggest new experiments that improve isoprenol production. By combining these tools we have shown how to pinpoint genetic modifications that improve production of isoprenol, a potential biofuel, by 23% (from 0.46 mMol to 0.57 mMol, Fig. 7), in a simulated data set. The same procedures are applicable in the case of real experimental data. In sum, this set of tools provides a solution for the data deluge that bioengineering is currently experiencing, and a way to build on preexisting data to fruitfully direct future research.

## Methods

### Synthetic data generator library (OMG)

The Omics Mock Generator (OMG) library is used to provide the synthetic multiomics data needed to test the computational tools described here (Fig. 2). Since experimental multiomics data is expensive and non-trivial to produce, OMG provides a quick and easy way to produce large amounts of multiomics data that are based on plausible metabolic assumptions. OMG creates fluxes based on Flux Balance Analysis and growth rate maximization (Orth et al., 2010), leveraging COBRApy (Ebrahim et al., 2013). OMG can use any genome-scale model, but in this case we have used the iJO1366 *E. coli* genome scale model, augmented with an isoprenol pathway obtained from the iMM904 *S. cerevisiae* model (Notebook A). In order to obtain proteomics data, we assume that the corresponding protein expression and gene transcription are linearly related to the fluxes. We also assume the concentration of metabolites to be loosely related to the fluxes of the reactions that consume or produce them (no fluxes → no metabolite): the amount of metabolite present is assumed to be proportional to the sum of absolute fluxes coming in and out of the metabolite. Therefore, although the data provided by OMG is not real, it is more realistic than randomly generated data, providing a useful resource to test the scaling of algorithms and computational tools.

The data generated by OMG was used in this manuscript to test EDD input, output, and visualization, and to provide training data for ART (Fig. 2). This tool can be a very useful resource for the rapid prototyping of new tools and algorithms.

#### Generating flux time series data

Fluxes describe the rates of metabolic reactions in a given organism, and can be easily generated through FBA. FBA assumes that the organism is under selective pressure to increase its growth rate (Orth et al., 2010; Lewis et al., 2012), hence searching for the fluxes that optimize it. FBA relies on GSMs, which provide a comprehensive description of all known genetically encoded metabolic reactions (Thiele and Palsson, 2010). FBA produces fluxes by solving through Linear Programming (LP) the following optimization problem:

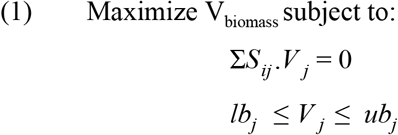

where *S* is the stoichiometry matrix of size *m*\**n* (number of metabolites*number of reactions in the model), *V_j_* is the flux for reaction *j* within the model (*j*=1,…,*n*). The lower bound (*lb_j_*) and upper bound (*ub_j_*) provide the minimum and maximum for each reaction. For example, if the input carbon source is glucose and we know that the input in a given time lapse is −15 mmol/gdw/h, the upper bound and lower bound for the exchange reaction corresponding to glucose are set to this value: −15 < V_EX_glc_ < −15. The solution to this optimization problem provides the fluxes that maximize growth rate, and that will be used later on to obtain transcriptomics, proteomics and metabolomics data.

We create time series of fluxes by doing a batch simulation based on FBA (see OMG library and Notebook A). We assume a given concentration for extracellular metabolites (e.g. 22 mM of glucose, or 18 mM of ammonium) and, for each time point, we run FBA for the model and update the extracellular metabolite concentration based on the exchange fluxes coming from the simulation (see Notebook A). For example, if an exchange flux of V_EX_glc_D = −15 mmol/gdw/hr is obtained for the model, the corresponding glucose concentration is adjusted as follows:

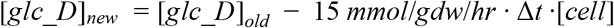

Where [*glc_D*]_*old*_ is the old glucose concentration, [*glc_D*]_*new*_ is the updated concentration, Δ*t* is the time change and [*cell*] is the cell concentration. In practice, we assume that the cell density increase is better described by an exponential than an linear relationship, and Δ*t* · [*cell*] is substituted by Δ[*cell*]/, where is the growth rate: = 1/[*cell*]·Δ[*cell*]/Δ*t*. The simulation proceeds until the carbon source (e.g. glucose) is exhausted. The result of the simulation is a set of fluxes, cell concentration and extracellular metabolite concentration for each time point (see Fig. 5). The fluxes will be the base for calculating transcriptomics, proteomics, and metabolomics data, as shown below.

#### Generating proteomics data

The flux values obtained from FBA are subsequently used to generate proteomics data, which describe the concentration of the protein catalyzing a given reaction within the host organism. The protein concentration for each time point is derived from the corresponding fluxes through a linear relationship:

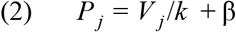

which is loosely inspired in the Michaelis-Menten equation (Heinrich and Schuster, 1996), where *V_j_* is the flux of reaction *j, P_j_* is the concentration of the protein catalyzing the reaction *j*, and *k* is a linear constant arbitrarily set to 0.1. The symbol *β* is an added random noise which is set to 5% of the signal.

#### Generating transcriptomics data

The aforementioned proteomics values are subsequently used to generate transcriptomics data, which describe the abundance of RNA transcripts linked to a given protein within the host organism. For simplicity, the transcript data is assumed to have a linear relationship with the proteomics data:

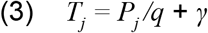

where *T_j_* is the abundance of RNA transcripts linked to the reaction *j*, and *q* is a linear constant arbitrarily set to 0.833. As above, *γ* is a random noise addition set to 5% of the signal data. This calculation is performed for each time point.

#### Generating metabolomics data

The flux values obtained from the FBA are also used to generate the metabolomics data, which describe the concentration of a given metabolite within the host organism. While finding the metabolite concentrations compatible with a given metabolic flux, protein concentrations, and transcript levels is a non-trivial endeavor, here we attempt to produce metabolite profiles that are not obviously unreasonable. Hence, we want concentrations of zero for metabolites that are connected to fluxes that are null, and non-zero in any other case. The easiest way to achieve this is by averaging the absolute value of all the fluxes producing or consuming the desired metabolite:

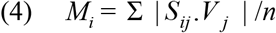

where *M_i_* is the concentration of the metabolite *i*, *j* is a reaction that involves metabolite *i*, and *n* is the total number of reactions in which metabolite *i* participates. This calculation is performed for each time point.

### Generating training data for machine learning and testing predictions

We leverage the OMG library to create training data to showcase the use of ART to guide bioengineering (Fig. 2). We will first create multiomics (transcriptomics, proteomics, metabolomics), cell concentration, and extracellular metabolite concentration data for the wild type *E. coli* strain (WT). Then we will use ART to suggest initial WT modifications (designs) so as to create enough data to train ART to be predictive. Those initial designs will be used by OMG to simulate isoprenol production data for bioengineered strains that include the genetic modifications indicated by the initial designs. ART will be trained on these data, and then used to suggest strain modifications that are predicted to increase isoprenol production. We will then compare ART predictions for isoprenol production with the “observed” results produced by using those designs to simulate bioengineered strains through OMG. In sum, OMG results are used as ground truth to be leveraged in testing ART’s performance.

This process involves the following phases:

1. **Choosing input and response variables.** Since the objective is to improve production of isoprenol, we use isoprenol concentration as the response variable. By inspecting the *E. coli* network we choose the following fluxes connected to acetyl-CoA, which is the source for the isoprenol pathway: ACCOAC, MDH, PTAr, CS, ACACT1r, PPC, PPCK, PFL. These fluxes then form the set of input variables for ART (Radivojević et al., 2020).
2. **Representation of different strain designs (i.e., genetic modifications).** We will consider only two types of modifications for each flux: knock-out (KO) and doubling the flux (UP). This choice results in three categories for each of the fluxes, which additionally include no modification (NoMod). We denote these categories by 0, 1 and 2 for KO, NoMod, and UP, respectively. Considering 8 fluxes and 3 options for design of each, the total number of possible designs is 3^8^ = 6561.
3. **Choose training data size**. We choose the initial training data to consist of 96 nonequivalent designs (instances), including the WT strain. This choice mimics one 96-well plate run. Different designs (instances) here represent different engineered strains. These 96 designs represent about 1.46% of all design space.
4. **Generate initial designs.** Initial designs are generated using ART’s feature for generating recommendations for the *initial* cycle, by setting its input parameter `initial_cycle` to True. This ART functionality relies on the Latin Hypercube method (McKay et al., 1979), which spaces out draws in a way that ensures the set of samples represents the variability of the full design space. Another parameter needed for ART is `num_recommendations`, which we set to 95 (see point 3 above). See the ART publication (Radivojević et al., 2020) for a list of other optional parameters. As the current version of ART deals only with continuous variables, ART’s recommendations will be drawn from interval [0, 1], which is the default interval if no specific upper and lower bounds are provided in a separate file. We then transform each of those values into one of the defined categories {0, 1, 2} by applying the function f(x) = 3*floor(x). Finally, we add a WT strain design {1,1,…,1}. See Jupyter Notebook B for the details.
5. **Generate production data for the initial designs.** The initial designs from ART are used as input to the OMG library, generating our “ground truth” for the isoprenol production levels for each of the initial designs. This represents the strain construction and the corresponding phenotyping experiments, which are simulated through OMG’s mechanistic modeling. In order to simulate how production is affected by the genetic changes suggested in the designs (e.g. knock out MDH, upregulate PFL and do not change CS), we used MOMA (Segrè et al., 2002) for each of the time points in the flux series (Fig. 10). Details can be found in Notebook C.
6. **Training ART with initial production data**. ART uses the initial designs and their corresponding productions from phases 4 and 5 to train, enabling it to predict isoprenol production for designs it has not seen. Details can be found in Jupyter Notebook D.
7. **Generate next-cycle design recommendations using ART**. Once trained, ART generates 10 recommendations that are expected to improve isoprenol production. Predictions are generated for each recommendation in the form of probability distributions (Fig. 7). Details can be found in Notebook D.
8. **Compare ART predictions to ground truth**. Finally, we take ART’s design recommendations from phase 7 and use OMG to simulate the corresponding ground truth isoprenol productions, similarly as in phase 5. We compare these isoprenol productions (observed production) versus the machine learning recommendations from ART (Fig. 8). Details can be found in Jupyter Notebook E.

**Figure 10:**
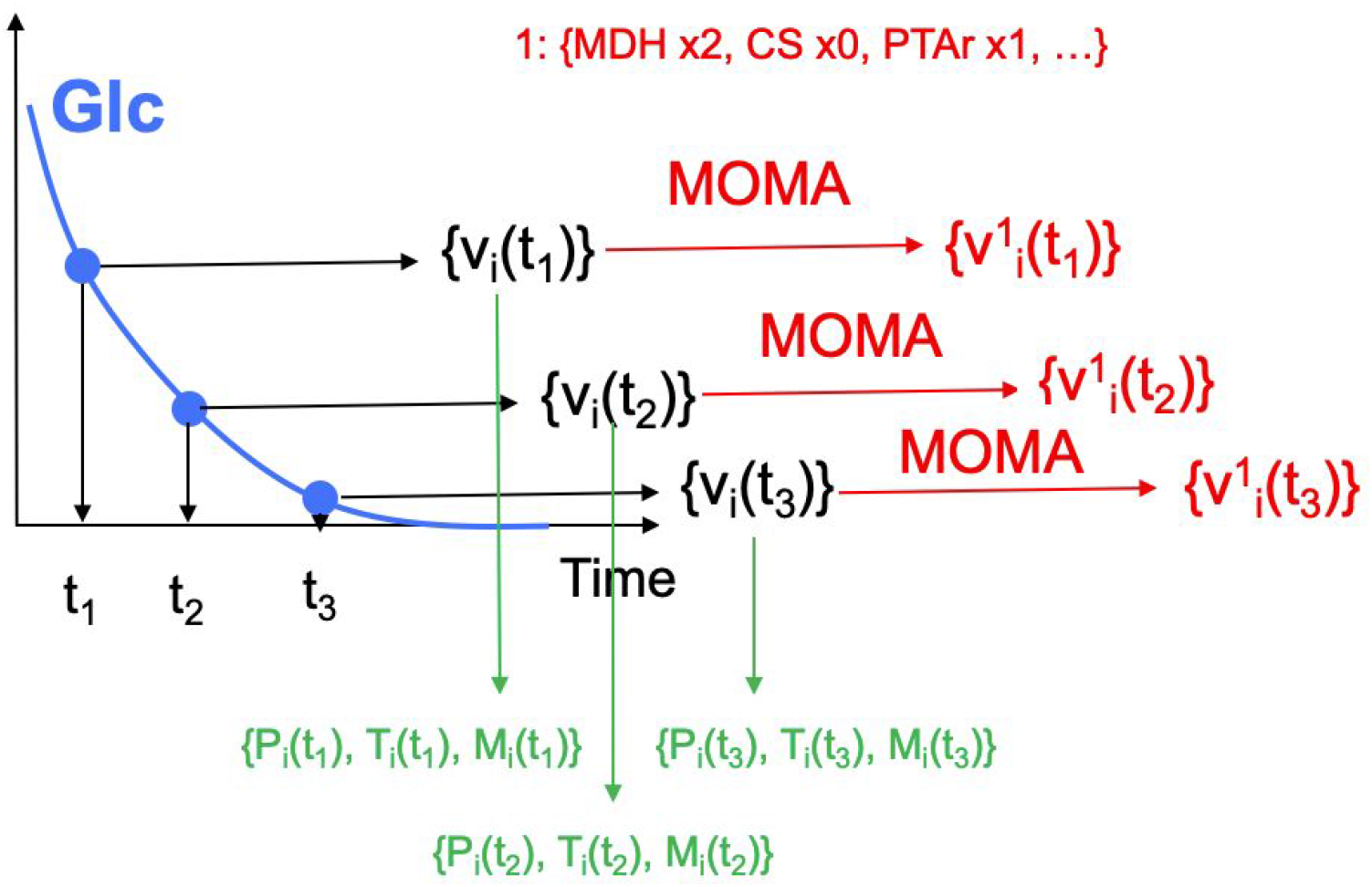
Generating multiomics time series data. For each time point, we generate fluxes by solving an FBA problem, until glucose is fully consumed. MOMA is used in conjunction with the *design* (e.g. increase CS flux two fold, knock POX out, maintain PFL) to predict fluxes for the strain bioengineered according to the design.

### Data formats for generic input files

The data formats for the generic input files in EDD involve five columns. The first one is the line it applies to (e.g. WT). The second one is the measurement type identifier, which involves a standardized choice: Pubchem IDs for metabolites, UniProt IDs for proteins, and Genbank gene IDs for transcripts (e.g., CID: 715, acetate). The third column is the time point (e.g. 0 hours). The fourth column is a value for the corresponding identifier (e.g. 2.4). And the final column is the unit corresponding to this value: FPKM for transcriptomics data, proteins/cell for proteomics data, and mg/L or mM for metabolomics data. Optical Density (OD) has no units.

## Code availability

All the code used in this paper can be found in the following github repositories:

- The Jupyter Notebooks associated with this paper, as well as the input files: https://github.com/AgileBioFoundry/multiomicspaper
- OMG: https://github.com/JBEI/OMG
- ICE: https://github.com/JBEI/ice
- EDD: https://github.com/JBEI/EDD
- The edd_utils package: https://github.com/JBEI/edd-utils
- ART: https://github.com/JBEI/ART

## Data availability

Data are available in the multiomics github repository. See Jupyter notebooks (Table 3) for exact file location.

## Author contributions

SR, TR, JMM, and HGM created and developed the concept for the paper. SR, JMM, TR and HGM developed the OMG library. TR and HGM created the Jupyter Notebooks. HGM created the screencasts. MF, WM developed the ART frontend. MF contributed error checking and bug fix code to ART. MF, NJH, TR and HGM designed and tested the ART frontend. SR, TR, MF, WM, JK, TB tested the screencasts and notebooks. SR, TR, MF, VJ, NH and HGM wrote the paper.

## Acknowledgements

This work was part of the DOE Agile BioFoundry (http://agilebiofoundry.org) and the DOE Joint BioEnergy Institute (http://www.jbei.org), supported by the U. S. Department of Energy, Energy Efficiency and Renewable Energy, Bioenergy Technologies Office, and the Office of Science, through contract DE-AC02-05CH11231 between Lawrence Berkeley National Laboratory and the U.S. Department of Energy. The views and opinions of the authors expressed herein do not necessarily state or reflect those of the United States Government or any agency thereof. Neither the United States Government nor any agency thereof, nor any of their employees, makes any warranty, expressed or implied, or assumes any legal liability or responsibility for the accuracy, completeness, or usefulness of any information, apparatus, product, or process disclosed, or represents that its use would not infringe privately owned rights. The United States Government retains and the publisher, by accepting the article for publication, acknowledges that the United States Government retains a nonexclusive, paid-up, irrevocable, worldwide license to publish or reproduce the published form of this manuscript, or allow others to do so, for United States Government purposes. The Department of Energy will provide public access to these results of federally sponsored research in accordance with the DOE Public Access Plan (http://energy.gov/downloads/doe-public-access-plan). This research is also supported by the Basque Government through the BERC 2018-2021 program and by Spanish Ministry of Economy and Competitiveness MINECO: BCAM Severo Ochoa excellence accreditation SEV-2017-0718.

The authors thank Zak Costello and Jacob Coble for their contributions during the initial development of the ART frontend.

## Supplementary material

The supplementary material is composed of a set of videos containing screencasts that showcase each part of the paper, and a set of Jupyter notebooks with all the code involved in the paper. These Jupyter notebooks allow anyone to replicate the procedures explained in the paper. These Jupyter notebooks and the input files they require can be found in the github repositories given above. Html versions of these notebooks, as well as the input files and the screencasts are provided in the SuppMaterial.zip file.

